# A performance-optimized model of neural responses across the ventral visual stream

**DOI:** 10.1101/036475

**Authors:** Darren Seibert, Daniel Yamins, Diego Ardila, Ha Hong, James J. DiCarlo, Justin L. Gardner

## Abstract

Human visual object recognition is subserved by a multitude of cortical areas. To make sense of this system, one line of research focused on response properties of primary visual cortex neurons and developed theoretical models of a set of canonical computations such as convolution, thresholding, exponentiating and normalization that could be hierarchically repeated to give rise to more complex representations. Another line or research focused on response properties of high-level visual cortex and linked these to semantic categories useful for object recognition. Here, we hypothesized that the panoply of visual representations in the human ventral stream may be understood as emergent properties of a system constrained both by simple canonical computations and by top-level, object recognition functionality in a single unified framework (Yamins et al., 2014; Khaligh-Razavi and Kriegeskorte, 2014; Güçlü and van Gerven, 2015). We built a deep convolutional neural network model optimized for object recognition and compared representations at various model levels using representational similarity analysis to human functional imaging responses elicited from viewing hundreds of image stimuli. Neural network layers developed representations that corresponded in a hierarchical consistent fashion to visual areas from V1 to LOC. This correspondence increased with optimization of the model’s recognition performance. These findings support a unified view of the ventral stream in which representations from the earliest to the latest stages can be understood as being built from basic computations inspired by modeling of early visual cortex shaped by optimization for high-level object-based performance constraints.

**Significance Statement:** Prior work has taken two complimentary approaches to understanding the cortical processes underlying our ability to visually recognize objects. One approach identified canonical computations from primary visual cortex that could be hierarchically repeated and give rise to complex representations. Another approach linked later visual area responses to semantic categories useful for object recognition. Here we combined both approaches by optimizing a deep convolution neural network based on canonical computations to preform object recognition. We found that this network developed hierarchically similar response properties to those of visual areas we measured using functional imaging. Thus, we show that object-based performance optimization results in predictive models that not only share similarity with late visual areas, but also intermediate and early visual areas.

## Introduction

Human cortex contains numerous areas with topographic representations of the visual world (Wandell et al., 2007; Silver and Kastner, 2009). What does each one of these cortical areas *do*? At least two major divergent approaches to this general question have been taken to understand areas in the ventral visual pathway which is thought to be involved in object vision and perception (Ungerleider and Mishkin, 1982; Goodale and Milner, 1992).

One approach, exemplified by research beginning with the primary visual cortex in cats (Hubel and Wiesel, 1959) and monkeys (Hubel and Wiesel, 1968), has been to examine the visual response properties of neurons and ask mechanistic questions about how properties such as orientation selectivity in simple cells and invariance to position in complex cells are created by neural circuitry (Hubel and Wiesel, 1962). This approach has led to computational models of visual cortical processing in which receptive fields are described as linear weightings (DeAngelis et al., 1993) of inputs from neurons with center-surround receptive fields (Kuffler, 1953; Hubel and Wiesel, 1961). As this linear weighting of visual inputs is performed by neurons with similar filtering properties tiled across the visual field, this stage of processing is akin to a convolution of a filter with visual input. Linear receptive fields are followed by output nonlinearites such as thresholding and exponentiation (Heeger, 1992; Gardner et al., 1999; Anzai et al., 1999) and divisive contrast normalization (Heeger, 1992). These basic computations are proposed to be canonical (Carandini and Heeger, 2012) such that repeating them in a hierarchical fashion (Fukushima, 1980; LeCun et al., 1990; Riesenhuber and Poggio, 1999) may recapitulate computations performed by visual areas along the visual pathways.

A second approach, exemplified particularly by research in humans (Kanwisher et al., 2001; Malach et al., 1995) and monkeys (Perrett et al., 1982; Desimone et al., 1984; Tanaka et al., 1991; Freiwald and Tsao, 2010; Hung et al., 2005) has started largely by asking about whether high-level features of visual scenes such as the presence of objects, faces, places and other identifiable semantic categories are represented in temporal cortex. Links between these representations and perception, for example with faces, are bolstered by similarities between the perceptual phenomenology (Tanaka and Farah, 1993) and representations in ventral cortex (Kanwisher and Yovel, 2006). Moreover, causal evidence in the form of lesion (Wada and Yamamoto, 2001) and stimulation evidence links high-level representations in the ventral visual stream in both monkeys (Afraz et al., 2006; Afraz et al., 2015) and humans (Parvizi et al., 2012) to perception.

Here we asked if a combination of these two approaches may help explain the nature of response properties not just of early and late areas, but for the full hierarchy of areas in human ventral visual cortex. We used a deep convolutional neural network model (Krizhevsky et al., 2012; Yamins et al., 2014) whose basic operations were inspired from the canonical computations derived from early visual cortex such as convolution, threshold non-linearities, non-linear pooling and normalization. We also constrained the network to develop high-level representations of object features, by training the network to perform well on invariant object recognition. Previous work has shown that these network models develop representational similarity to V4 and IT in monkey (Yamins et al., 2014; Khaligh-Razavi and Kriegeskorte, 2014) and humans (Khaligh-Razavi and Kriegeskorte, 2014; Güçlü and van Gerven, 2015). We capitalized on the ability to measure responses in multiple topographically and functionally localized cortical areas of the human using BOLD imaging to see if this framework could be extended to the whole ventral stream from earliest cortical stages to later ventral areas. While intermediate visual areas such as V2 might be expected to have some kind of intermediate representation between V1 and later stages of the visual system (there are many possible such representations), our model was not explicitly trained to fit V2 responses and therefore was not guaranteed to show any correspondence. Nonetheless, we found representational similarity between the neural network and the human visual system in a hierarchical consistent fashion.

## Materials and Methods

### Human subjects

Seven subjects (1 female, ages 22-38) participated. Subjects provided written and oral informed consent before each session and all procedures were approved by the Author University. All subjects underwent at least four imaging sessions (anatomical, retinotopy, category localizer and main experiment). Similar to other studies (Kay et al., 2008; Naselaris et al., 2009; Stansbury et al., 2013), our analyses required consistent responses to hundreds of image stimuli over many scanning sessions from each subject. Therefore, from the original cohort of subjects, we selected the two which had the highest mean split-half reliability in V1 (see the Stimulus response section) to complete a full data set (at least 9 sessions each consisting of approximately 10 8-minute scans of the main experiment). Of the two pre-screened subjects chosen to complete the full dataset, one was an author. This pre-screening procedure was designed to select subjects based on the overall reliability of data without introducing bias for what representations subjects exhibit. We note that because of the design decision to collect a large data set from a small number of subjects, the results presented here are generalizable only if visual representations in the ventral visual areas across individuals is similar - a notion that is supported by a great deal of literature both within and across species of primates (Kriegeskorte et al., 2008; Yamins et al., 2014; Khaligh-Razavi and Kriegeskorte, 2014; Kay et al., 2008; Naselaris et al., 2009; Stansbury et al., 2013; Güçlü and van Gerven, 2015).

### Stimuli

We presented 1785 gray-scale images of objects a median of six times across multiple sessions to each subject. Objects were drawn from 8 categories (animals, tables, boats, cars, chairs, fruits, planes, and faces) containing 8 exemplars. Each object was shown from 27 or 28 different viewpoints against a random natural background (circular vignette, radius 8° centered on fixation) to increase object recognition difficulty (Figure 1). We used a rapid event-related design where each image was presented for 1.25 s followed by a random delay between 1 and 4 s. Subjects maintained fixation while performing a 2AFC luminance decrement discrimination task on the fixation cross (Gardner et al., 2008) whose timing was randomly out of sync with stimulus presentation.

**Figure 1.**
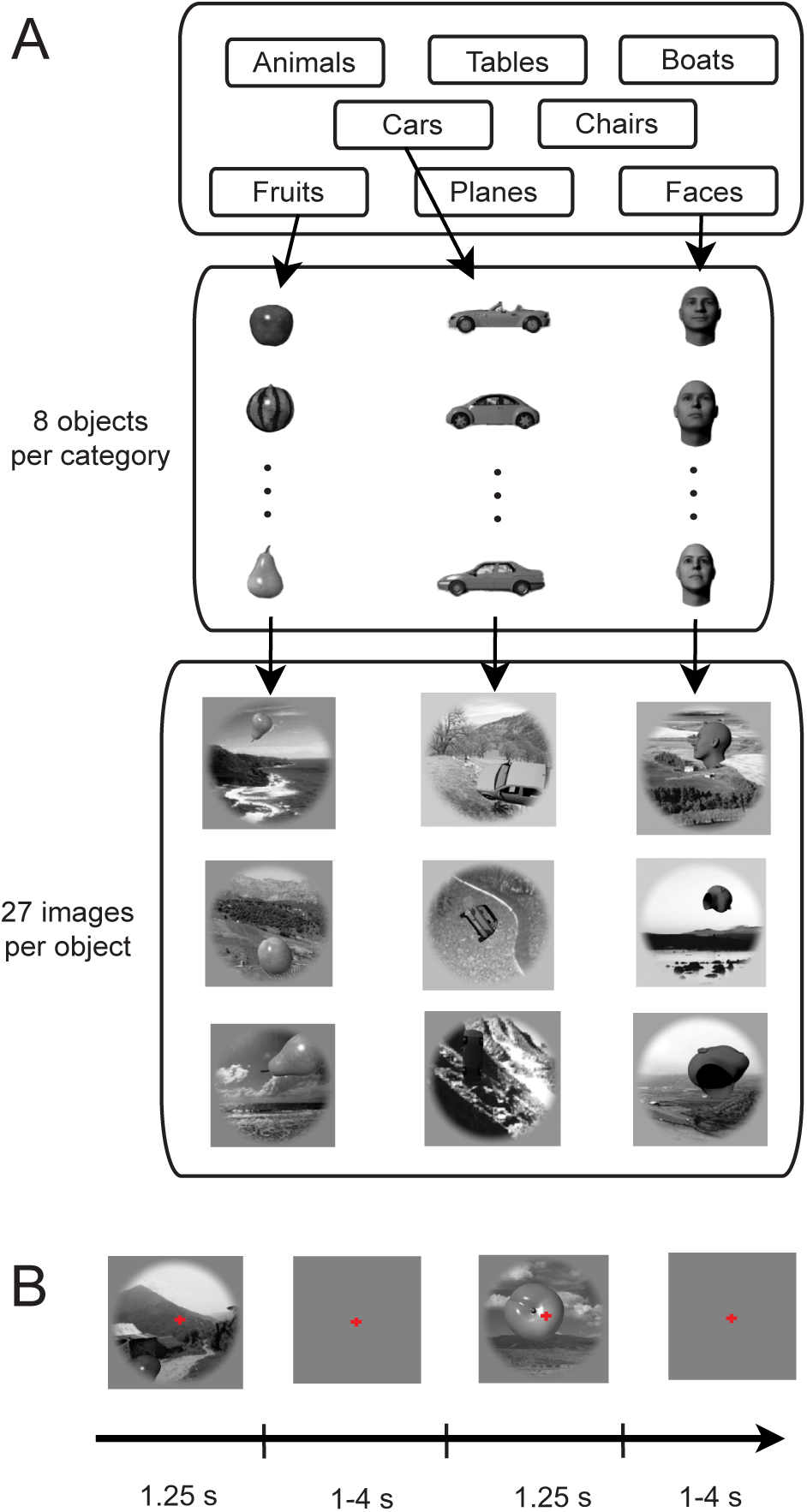
Task and stimuli. **A**, Stimuli contained 8 objects chosen from 8 categories. Each object appeared in 27 or 28 images in random positions, scales, orientations, and on random backgrounds. **B**, Images were shown for 1.25 s followed by a random delay of 1-4 s. Subjects maintained central fixation and performed a discrimination task on the fixation cross.

### MRI methods

Data were collected at the Author University with a Varian Unity Inova 4T whole-body MRI scanner using a head gradient system (Agilent). We collected a T1-weighted anatomical scan (MPRAGE; TR, 13 ms; TI, 500 ms; TE, 7 ms; flip angle, 11°; voxel size, 1 × 1 × 1 mm; matrix, 256 × 256 × 180) and a T2-weighted anatomical images (TR 13 ms, TE 7 ms, flip angle 11°, matrix 256 × 256 × 180; 1 mm isotropic voxels) for each subject. We divided the T1 and T2-weighted images to correct for contrast inhomogeneities (Van de Moortele et al., 2009) and segmented this reference anatomical to generate cortical surfaces using Freesurfer (Dale et al., 1999).

We collected functional scans at 3 × 3 × 3 mm (matrix size, 64 × 64 × 27) using echo-planar imaging. Scans were collected with a TR of 1.25 s, a TE of 25 ms, flip angle 30^°^ using sensitivity encoding (acceleration factor of 2) (Pruessmann et al., 1999). We showed 210 distinct images each session (105 stimuli per run, alternating between two run types). In each functional session, we collected an anatomical scan for cross-session alignment to each subject’s high-resolution anatomical.

Subject 1 (S1) participated in 14 functional sessions and was shown 2539 images. Subject 2 participated in 9 functional sessions and shown 1785 images–a subset of those shown to S1. Our analyses used the 1785 images shown to both subjects.

### MRI data preprocessing

We recorded physiological data to reduce noise artifacts. Respiration measurements from a pressure sensor and pulse oximeter data were used for retrospective estimation and correction in *k* space (Hu et al., 1995). tSENSE (Kellman et al., 2001) acceleration artifacts were removed with notch filtering using mrTools. No slice time correction or spatial smoothing was performed. We corrected head motion using standard approaches (Nestares and Heeger, 2000).

### Visual area definitions

We collected one retinotopic session for each subject (Gardner et al., 2008; Wandell and Winawer, 2011). The imaging parameters were the same as our functional sessions (exceptions: *r* = 2, tSENSE acceleration, effective TR 1.02 s, 35 axial slices). We positioned slices perpendicular to the calcarine sulcus. Preferred angle and eccentricity for each voxel were estimated using a Fourier-based analysis and projected on the gray matter surface.

We used 6 runs for our retintopic area definitions. Two runs of both clockwise and counterclockwise wedges were used and one run each of expanding and contracting rings. In each run we collected 168 volumes (24 volumes per cycle, 10.5 cycles). We discarded the first half cycle to minimize visual adaptation effects. While maintaining fixation, subjects performed a staircased two-alternative forced choice contrast discrimination task at fixation to maintain alertness.

Similar preprocessing was performed on the retinotopic sessions as the main experiment. After preprocessing, we time reversed (2 volume offset to correct for hemodynamic lag and improve SNR) the counter-clockwise runs and averaged together these runs with the clockwise runs. This left us with an average time-series for the ring and wedge runs. We determined the preferred angle and eccentricity phase for each voxel using a Fourier-based correlation analysis. We projected these values on the flattened gray matter surface and defined border definitions using published procedures (Wandell et al., 2007). V1, V2, V3, V3A, hV4, LO1, and LO2 were defined in the ventral stream (Schluppeck et al., 2005; Swisher et al., 2007; Silver and Kastner, 2009).

### Category area definitions

Imaging parameters for the category localizer session were the same as functional sessions (exceptions: *r* = 4, tSENSE acceleration, effective TR 1.08 s). We showed natural images matched to have identical magnitude in Fourier space to reduce differences between object categories (Rajimehr et al., 2011). Scrambled and intact images were shown at 14° height and width. The session was block designed with 12.9 s blocks, 13 images per block 0.75 s on, 0.25 s off.

Preprocessing for the localizer session was similar to that of our main experiment; however we applied spatial Gaussian smoothing (6 mm full width at half maximum). We created a design matrix with predictors for each of the block types by convolution with a canonical hemodynamic response function (difference of gamma functions, *x* = 6, *y* = 16, *z* = 6, where *x* and *y* were the shape parameters of the positive and negative functions and *z* was the ratio of the scaling parameter of positive to negative gamma functions). Using the design matrix, we fit a GLM to each subject’s data individually. Using the fitted responses, we calculated a contrast for intact stimuli (scenes, faces, and natural objects) to scrambled. We defined and masked LOC using a statistical threshold of *p* ≤ 0.0001 (uncorrected) and removed all voxels within the retinotpically defined areas (V1-V4). We defined PPA, OFA, and FFA using similar procedures. OFA and FFA were defined using a faces to objects contrast (Kanwisher et al., 1997). PPA was defined using a scenes over objects contrast (Epstein and Kanwisher, 1998). Some of LOC overlapped with LO2, however it should be mentioned LOC is not a superset of LO1 plus LO2, as they are defined using entirely separate criteria (category localization versus retinotopy) (Larsson and Heeger, 2006).

### Image responses

We used GLMs with PCA components of non-visually driven voxels as noise regressors to estimate image responses of each voxel with GLMdenoise using the package’s default HRF (Kay et al., 2013), which produced for each voxel one response (GLM coefficient) for all presentations (median of 6) of each image. We computed reliability by randomly splitting the scans into two groups and estimating responses for each group. The correlation between the vectors was our estimate of split-half reliability. We discarded voxels with *r* ≤ 0 (similar to Mitchell et al. (2008)) and pooled voxels across subjects resulting in 536 voxels for V1, 407 for V2, 510 for V3, 379 for hV4, 123 for PPA, 192 for OFA, 292 for FFA, 234 for LO1, 299 for LO2, 111 for TOS, and 535 for LOC. Our analyses are based on the assumption that ventral visual representations are similar across subjects, based on prior work which has shown remarkable representational similarity not only across subjects but across species (Kriegeskorte et al., 2008; Khaligh-Razavi and Kriegeskorte, 2014).

### Convolutional neural network architecture

We used a convolutional neural network (CNN) inspired by Krizhevsky et al. (2012). Our model consisted of two branches of three main layers. Each main layer contained one or more convolutional stages followed by normalization and pooling. Figure 5 illustrates the architecture of our network. Normalization and pooling followed the first, second, and fifth convolutional stages. We used the publicly available cuda-convnet package with minor custom modifications to train and evaluate our model (Krizhevsky et al., 2012). Our main analyses focus principally on the outputs of the three main layers.

Each of the 5 convolutional stage was constructed using rectified linear units. Rectified linear units are a simple non-linearity of the form *f*(*x*) = *max*(0, *x*) and were chosen by Krizhevsky et al. (2012) in part because training networks with this form of non-linearity is quicker 209 than other non-linearities. The five convolutional stages contained filters of spatial sizes 11 × 11, 5 × 5, 3 × 3, 3 × 3, and 3 × 3 px. Each convolutional stage had 48, 128, 192, 192, and 192 filters respectively.

We used 3 identical response normalization stages as Krizhevsky et al. (2012). For a given unit *a^i^_x,y_* representing the activation of channel *i* at spatial position *x, y*, the normalized output is defined as,

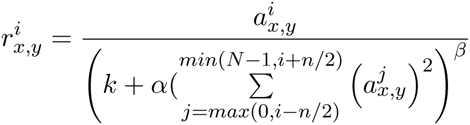

where *n* is the number of channels in the same spatial location to normalize across, and *N* is the number of channels in the layer. Because we initialize all convolutional weights randomly, the ordering of the channels is initially arbitrary. Like Krizhevsky et al. (2012), we set *k* = 2, *n* = 5, α = 10^−4^, *β* = 0.75.

Our 3 max pooling stages were also defined as in Krizhevsky et al. (2012). Max pooling takes the maximum value across space in each channel. We used max pool windows of size 3 × 3 with a spatial distance of 2 units between each pooling window. Using a smaller distance between windows than the size of the windows results in overlapping pooling, which Krizhevsky et al. (2012) observed results in a modest boost in model performance than non-overlapping pooling.

We simplified the architecture described by Krizhevsky et al. (2012) based on a preliminary analysis of which aspects of the model influenced performance on the 2013 ImageNet challenge-set. Namely, we removed two of the middle fully connected layers (compromising the majority of the model’s free parameters). Because the only remaining fully connected layer in our model was the top-layer (the classifier outputs), we did not utilize drop-out, unlike Krizhevsky et al. (2012). We additionally reduced the overall size of the network by reducing the input image size from 224 × 224 px to 120 × 120 px. With the training/test split of the 2013 ImageNet challenge-set we observed no significant changes in model performance after making these changes.

### Convolutional neural network optimization

The fitting procedure used here follows that of Krizhevsky et al. (2012). We learned filters and bias terms for each convolutional stage and the final fully connected layer with stochastic gradient descent. Batch sizes of 128 images were used from the 2013 ImageNet challengeset. The model was not trained on any synthetic images. All normalization and pooling parameters were held fixed and chosen to match Krizhevsky et al. (2012). In total, 9,019,111 parameters were learned. The majority of these parameters (6,912 × 999 = 6,905,088) were weights for the fully connected layer, which can essentially be thought of as classifier weights for the ImageNet challengeset–the output of the fully connected layer (a vector of 999 elements) is directly normalized to give the probability that a given image belongs to each of the 999 categories. Excluding the fully connected layer, which we did not use in subsequent analyses, the remaining five convolutional stages contributed 3 × 11 × 11 × 48, 48 × 5 × 5 × 128, 128 × 3 × 3 × 192, 192 × 3 × 3 × 192, and 192 × 3 × 3 × 192 weighting parameters respectively per branch, in addition to 48, 128, 192, 192, and 192 bias parameters per branch, for a total of 2,113,024 parameters.

Backpropagation training was performed for several days on a single NVIDIA Titan GPU for 74 epochs. To prevent overfitting, we augmented the training set by randomly cropping 120 × 120 px image patches from re-scaled 130 × 130 px images of the 2013 ImageNet challenge-set. Weights were initiated from a zero-mean Gaussian distribution with a standard deviation of 0.01. We manually reduced the learning rate of the procedure an order of magnitude when we observed the log-probability on the testing-set no longer decreased. Three such reductions in learning rate were performed. We terminated the fitting procedure upon observing further reductions in learning rate did not produce any additional decrements in the log-probability. The final performance value of the model reached that of ~70% correct (chance = 0.1% correct) and was within error of Krizhevsky et al. (2012).

### Control models

We included three controls: V1-like (Pinto et al., 2008), V2-like (Freeman and Simoncelli, 2011), and HMAX (Serre et al., 2007) models. V1-like consisted of Gabor filters at multiple scales, orientations, phases, and frequencies. V2-like consisted of non-linear conjunctions of Gabor outputs. HMAX contained hierarchical operations inspired by V1. We included an animate-inanimate RDM, created on the categorical animacy of each stimulus. The animate-inanimate RDM represents something of an upper bound to which increased categorization performance can lead to increased representational similarity for higher visual areas.

The HMAX model was built on similar principles to our CNN. It contained linear-non-linear layers involving filtering and max poolings. The architecture and training procedure of HMAX and our CNN, however differ. HMAX, for instance, contains approximately an order of magnitude less trainable parameters (10^5^ vs 10^6^) and is a shallower architecture. In addition, its training procedure is not gradient-based, making it somewhat less optimal in any given training regime. These properties make HMAX a reasonable intermediate control between our V1-like control model and our CNN. To give the HMAX model the best possible chance to perform, we pretrained the model using the stimulus images used to evaluate the model (for which we have BOLD data). This is in contrast to our CNN which was never trained on any images shown to our human subjects (or even any synthetic, 3D generated images).

### Representational dissimilarity matrices

We computed representational dissimilarity matrices (RDMs), like Kriegeskorte et al. (2008), consisting of one minus the pair-wise correlation of feature vectors (where features were GLM coefficients for each voxel in the case of brain areas and model unit outputs in the case of the model). Diagonal entries were set to 0.

Compared to other studies (Kriegeskorte et al., 2008; Khaligh-Razavi and Kriegeskorte, 2014), we used a far larger stimulus set where each object appears in multiple images shown in different positions, orientations, and scales. Because we were interested in the emergence of object perception, we created RDMs of object-averaged response vectors where we average features across images representing the same object. The object-averaged RDMs were also necessary to increase the amount of signal in our data–our stimulus set was purposely designed to be very difficult for observers to recognize the objects in order to expose the key computational aspects of invariant object recognition. Even when given infinite viewing time, there are many images in our stimulus set that human observers cannot recognize due to extreme variations in pose, orientation, and scale.

Because responses in each imaging voxel likely result from the activity of multiple neurons with different feature selectivities, we used a linear re-mapping of model features (c.f. Khaligh-Razavi and Kriegeskorte (2014)). We computed the correlation between model layers RDMs and visual response RDMs using a linear re-mapping of model features to match a given visual area’s RDM–each model layer and visual area pairing had its own set of weightings. The advantage of this approach is that it does not require model features be precisely synonymous with voxels which reflect large collections of neurons with potentially varying selectivities. The disadvantage of re-mapping is that it may be prone to overfitting, which we address with cross-validated bootstrapping and regularization. To estimate effect sizes, we used cross-validated bootstrapping which has the advantage of estimating our fitting reliability but is disadvantageous in that it requires us to fit on random subsets of the dataset rather than all of it. Each training set consisted of 1000 randomly selected model outputs to 15 images for each of 64 objects (960 images total). Model outputs for the remaining 12 images per object were used for testing. We found the vector *w* that maximizes *corr*(*RDM*(*V*), *RDM*(*X* ○ *W*)), where *corr*() is the Pearson correlation, *RDM*() is the vector of pair-wise row correlations, *V* is the matrix of object-averaged voxel responses (objects by voxels), *X* is the matrix of object-averaged model features (objects by 1000), *W* (objects by 1000) consists of rows of *w*, and ○ represents point-wise multiplication. We find *w* using the L-BFGS-B algorithm (Byrd et al., 1995) for 1 iteration (to both reduce computational time and as a form of early stopping to prevent overfitting). We report the average correlation on the testing set over 100 bootstraps (Figure 2) and 10 bootstraps (Figure 4). We used this procedure to calculate correlation values for all model layers as well as for all control models. Our linear re-weighting procedure is closely related but not identical to Khaligh-Razavi and Kriegeskorte (2014). Khaligh-Razavi and Kriegeskorte (2014) fit one weight per layer or model instead of per feature. With the correct normalization, squared Euclidean distances are proportional to correlation distances and non-negative least-squares on this quantity should maximize the RDM correlation distance like the method we used here.

**Figure 2.**
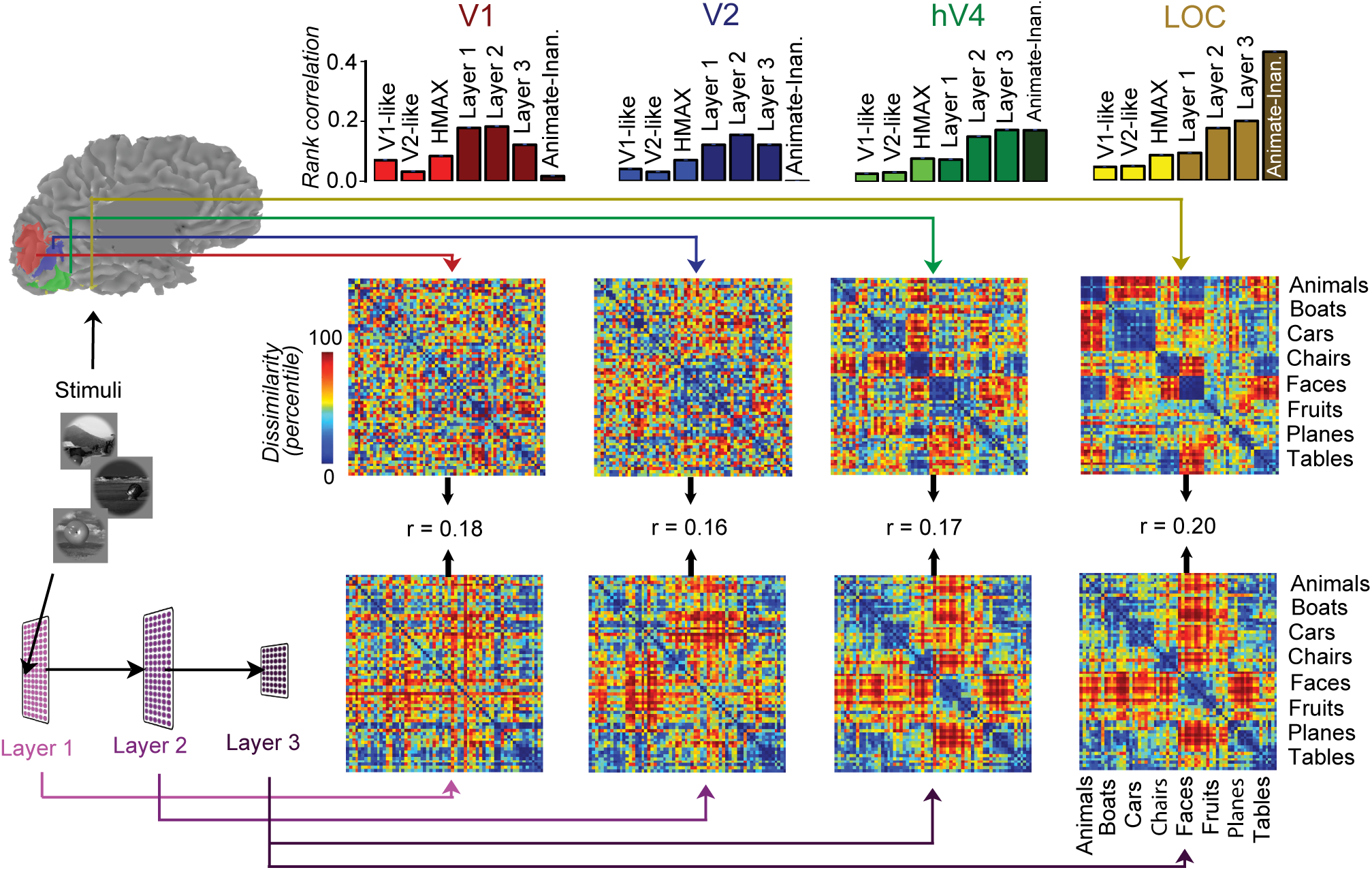
Model and neural representations. **Left**, Human functional imaging data and model responses to the same stimuli were used to compute RDMs at different levels of the visual system (top row) or layers of the model (bottom row). Increasing block-diagonality of the RDMs from V1 to LOC and from Layer 1 to Layer 3 illustrate emergence of categorical representations. Rank correlations between model layers and visual area RDMs **Top**, showed better correspondence than control models (V1-like, V2-like, and HMAX). Bars indicate SEM over bootstraps (see Materials and Methods).

**Figure 3.**
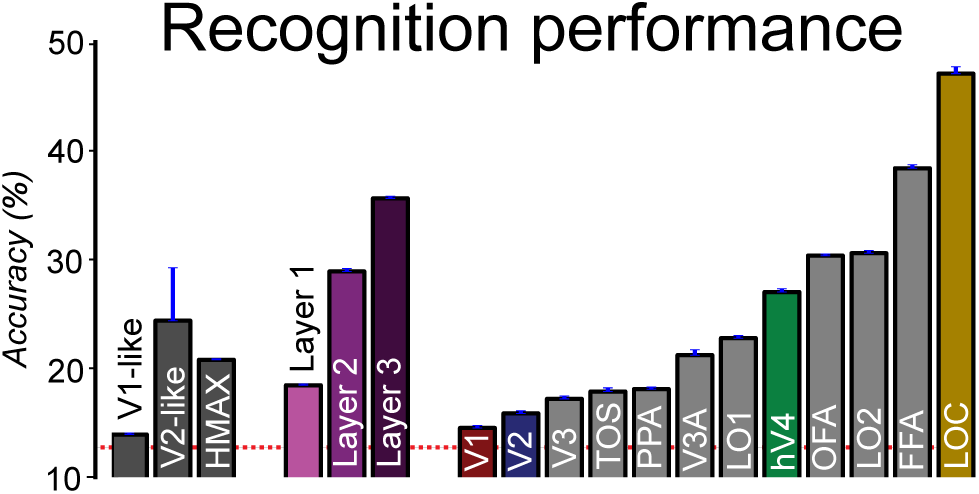
Object recognition performance. Accuracy for each model and visual area was computed with a cross-validated linear support vector machine (chance = 12.5%; dashed red line). The same training/test procedure was used for model and neural responses.

**Figure 4.**
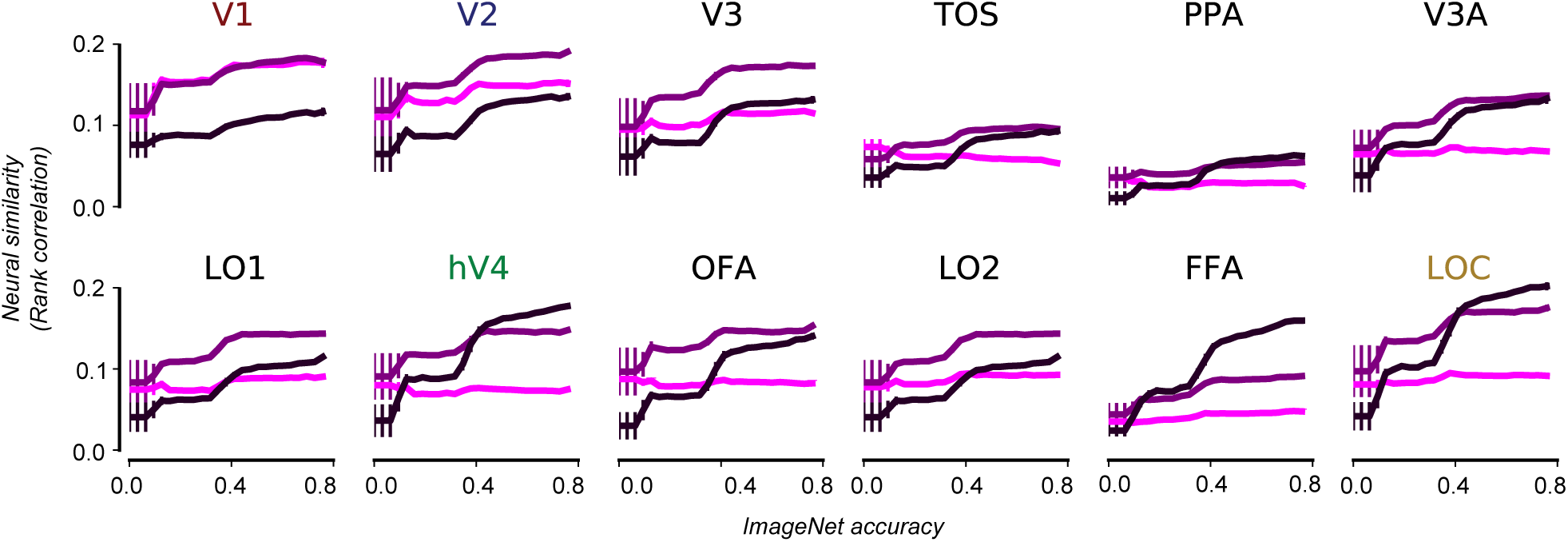
Optimization at object recognition performance improved model and visual area correspondence. Correlations between RDMs of each model layer (different colors; light purple: Layer 1, medium: Layer 2, dark: Layer 3) and visual area (different graphs) are shown as a function of model performance on ImageNet taken at different “checkpoints”. A positive trend indicates that, as the model becomes optimized on ImageNet recognition, it is better able to explain neural responses. Vertical bars indicate SEM over checkpoints (they become eclipsed by the width of the line plot on the far right of the plots).

We were not able to reliably calculate split-half explainable variance estimates for this linear re-mapping procedure due to the difficulty of fitting weights on smaller fractions of our data. However, these estimates were not critical to the hypotheses tested in this study because we were comparing the relative ranking of model predictivity for each visual area (ex. layer *X* explains visual area *A* significantly more than layer *Y*). To avoid the problem of finding linear re-weightings using smaller sub-sets of our data, we instead computed noise ceilings and percent explained variance values (Figure 6) without using the weighting procedure described above. Noise ceilings for each visual area were computed by splitting the runs of our data into two non-overlapping groups. With each group, we estimated stimulus responses (beta weights) using the procedure described above (see the Image responses section) and computed object-averaged RDMs for each visual area. We used the correlation between the RDM from each of the two groups as our noise ceiling for percent explained variance estimates (Figure 6).

**Figure 5.**
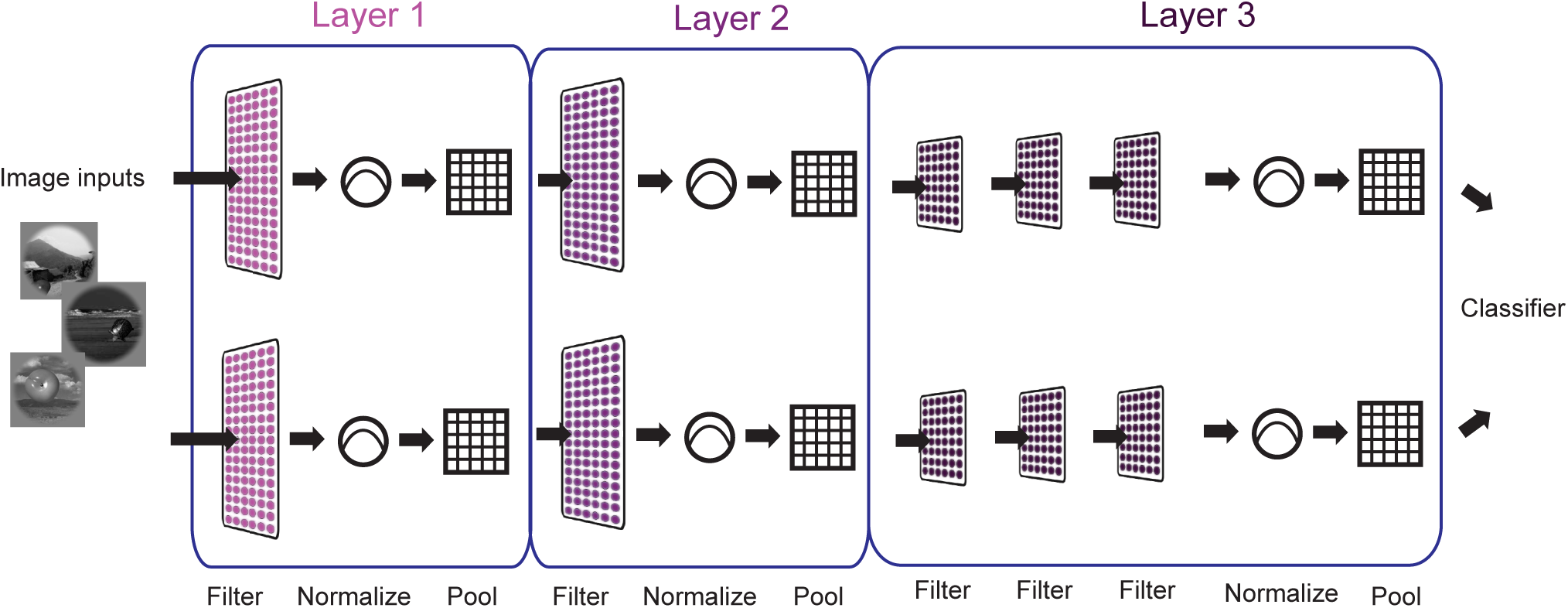
Architecture of our convolutional neural network. Beginning with the first layer and extending through until the fully connected (classifier) layer, the model contains two branches. The first convolutional layers for both branches each contain 48 filters, followed by 128 filters in the second convolutional layer, 192, 192, and 192 filters for the third, fourth, and fifth convolutional layers. We used the same filter sizes (11 × 11, 5 × 5, 3 × 3, 3 × 3, and 3 × 3 px) for convolutional layers 1-5 and striding parameters as Krizhevsky et al. (2012).

**Figure 6.**
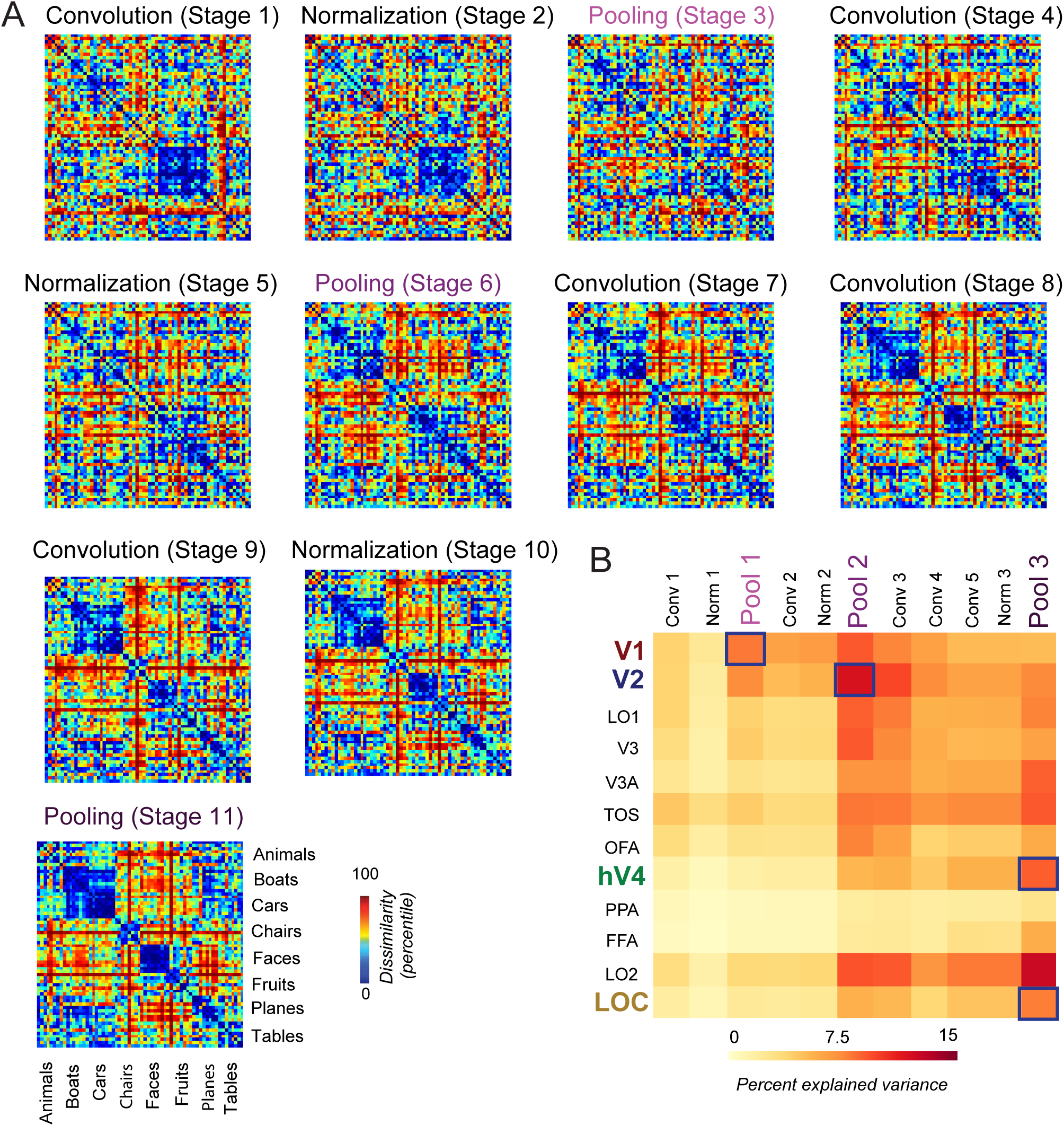
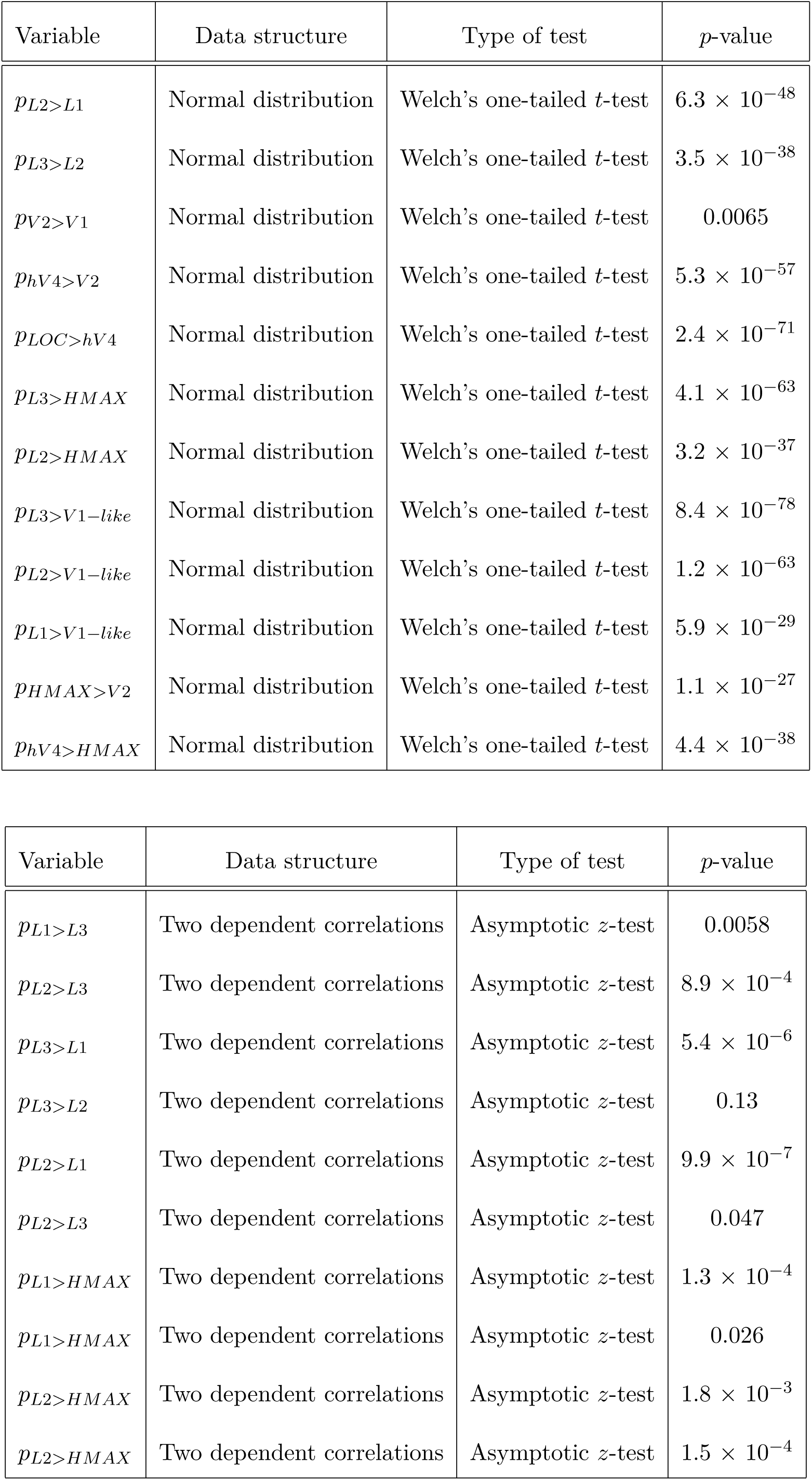
Object-averaged RDMs for all model stages. **A**, Shown are the unweighted model RDMs–they are not re-weighted to any of the fMRI visual area responses, unlike Figure 2. The pooling stages represent pooling layers 1 through 3 which were used in our main analyses. **B**, Shown are the percent explained variances between each model stage and visual area. We did not re-weight the model features in this analysis–instead of sampling 1000 random features we used all features from a given model stage. Blue boxes indicate, for each selected ROI, which model layer exhibited the highest correlation to the ROI.

### RDM statistical analysis

Using the bootstrapping above, we computed *p*-values testing if Layer *A* better explained visual area *X*’s RDM than Layer *B* (where *A* = 1 and *B* = 3, *X* =V4, for example). We use the notation *p_LA<LB_* to denote the *p*-value of *r_V, A_* < *r_V, B_*, where *r_V, A_* is the testing-set Spearman correlation of layer *A* and visual area *V*’s RDMs averaged over bootstraps. We use Fisher’s *r*-to-*z* transformation using Steiger (1980)’s approach to compute *p*-values for difference in correlation values (Lee and Preacher, 2013). The approach tests for equality of two correlation values from the same sample where one variable is held in common between the two coefficients (in our case, an RDM of a given visual area). We report p-values which are not corrected for multiple comparisons. Our approach bootstraps over independent stimulus samples and avoids problems that can arise from randomly sampling RDM matrices directly. Direct sampling of the RDM (ex. randomly sampling elements from it) can be problematic because two such random samples are not independent–a single stimulus contributes to multiple elements in the RDM matrix (Nili et al., 2014).

Spearman rank correlations are known to be biased for RDMs containing many tied ranks and can produce artificially high correlations (Nili et al., 2014). While the animate-inanimate control RDM has many tied rankings, none of our model or visual area RDMs contain tied ranks. For this reason, using Spearman correlations with the animate-inanimate RDM may produce misleadingly high correlations, particularly for higher visual areas. As noted in Nili et al. (2014), Kendall’s Tau correlation penalizes tied ranks, however, empirically, for our data–set it does not produce qualitatively different results. That is, even with Kendall’s Tau correlation, the animate-inanimate RDM significantly out-performs all model layers (ex. for a single bootstrap we observe a Tau value of 0.328 for animate-inanimate to LOC vs. a Tau value of 0.121 for layer 3 to LOC).

### Classification

We assessed model and neural recognition performance with cross-validated linear support vector machines (SVMs). Classifiers were trained on stimulus category of individual image responses. Training consisted of 20 random presentations of each object and testing consisted of the remaining presentations. We report median accuracy over 20 bootstraps. We set the classifier regularization “C” parameter equal to 0.0005 and computed significance by a one-tailed Welch’s *t*-test. We have not performed corrections for multiple comparisons.

### Performance vs. fitting

During ImageNet optimization, we measured model and neural similarities. At 100 gradient updates (checkpoints) spaced evenly through optimization, we computed RDM correlations using the procedure above. We sampled 100 points spaced evenly over the range of model performance values and plotted the average correlation over model checkpoints within 0.10 accuracy of each sampled point.

## Results

We optimized a convolutional neural network model for object recognition on a challenging image-set (Deng et al., 2009) to test the extent it matched the human visual system. After optimizing using backpropagation, the model achieved ~70% accuracy (chance = 0.1%) on ImageNet, and comparable although slightly reduced performance relative to humans, consistent with previous work (Krizhevsky et al., 2012; Yamins et al., 2014).

Emergence of categorical information was evident in model and human representations. We computed object-averaged RDMs (Kriegeskorte et al., 2008) for visual areas and model layers (see Materials and Methods). Each entry in an RDM is a measure of how dissimilarly a pair of objects are represented. Arranging the stimuli by category, we observe the emergence of block-diagonality (Figure 2). Blocks correspond to the emergence of categorical tolerance through the ventral stream, as within-category similarities are increasingly abstracted despite the high levels of variation in the stimuli. The RDMs of the model (Figure 2) also evidence emergence of categorical information.

We quantified recognition performance in model and visual areas by training support vector machines (SVM) to decode the category of each stimulus response (Figure 3). We observe increasing performance as we move from lower to higher model layers (*p*_*L*2>*L*1_ = 6.3 × 10^−48^; *p*_*L*3>*L*2_ = 3.5 × 10^−38^; see Materials and Methods: Classification) and increased performance as we move from posterior to anterior areas (as shown in Figure 3; *p*_*V*2>*V*1_ = 0.0065; *p*_*hV*4>*V*2_ = 5.3 × 10^-57^; *p*_*LOC*>*hv*4_ = 2.4 × 10^−71^). V1-like and HMAX models generally perform worse than the layers of our model (*p*_*L*3>*HMAX*_ = 4.1 × 10^−63^; *p*_*L*2>*HMAX*_ = 3.2 × 10^−37^). V1-like performs similarly to the fMRI V1 responses (but worse than all of our model layers-*p*_*L*3>*V*1–*like*_ = 8.4 × 10^–78^; *p*_*L*2>*V*1–*like*_ = 1.2 × 10^−63^; *p*_*L*1>*V*1-*like*_ = 5.9 × 10^−29^). HMAX performs in between V2 and hV4 responses (*p*_*HMAX*>*V*2_ = 1.1 × 10^−27^; *p*_*hV*4>*HMAX*_ = 4.4 × 10^−38^).

We found correspondence between model pooling layers and visual areas. Early areas were best explained by early layers and later areas by later layers (Figure 2A, e.g. compare layer correlations of V1 to LOC). V1 was best explained by lower–layers (*p*_*L*1>*L*3_ = 0.0058; *p*_*L*2>*L*3_ = 8.9 × 10^−4^; see Materials and Methods: RDM statistical analysis), and LOC was best explained by higher layers (*p*_*L*3>*L*1_ = 5.4 × 10^−6^; *p*_*L*3>*L*2_ = 0.13; *p*_*L*2>*L*1_ = 9.9 × 10^−7^). We observed intermediate visual areas, such as V2 and hV4, following this trend. V2, for instance, was better explained by the middle Layer 2 than the top layer (*p*_*L*2>*L*3_ = 0.047).

Our model exhibited higher similarity to the ventral stream than several control models: a V1-like model (Pinto et al., 2008), a V2-like model (Freeman and Simoncelli, 2011), and HMAX (Serre et al., 2007) (ex. for V1 *p*_*L*1>*HMAX*_ = 1.3 × 10^−4^; for V2 *p*_*L*1>*HMAX*_ = 0.026, for hV4 *p*_*L*2>*HMAX*_ = 1.8 × 10^−3^, and for LOC *p*_*L*2>*HMAX*_ = 1.5 × 10^−4^; see Materials and Methods: RDM statistical analysis). HMAX, V1-like, and V2-like models predicted hV4 and LOC RDMs approximately as well as Layer 1 of our model. For earlier visual areas, the control models were significantly worse at predicting the neural RDMs than any layer of our model (see aforementioned statistics). Our model exhibited lower correlations than the animate-inanimate RDM in LOC. However, unlike other controls, the animate-inanimate RDM does not represent the outputs of an image-computable model. The animate-inanimate RDM represents something of an upper bound in terms of how far we might expect increased performance optimization to lead to increased neural fitting of higher visual areas. It should be noted that we have not arranged the rows and columns of our RDMs in a way that visually highlights the animate-inanimate distinction observed previously (Kriegeskorte et al., 2008). However, the animate-inanimate RDM correlations are a quantitative measure of this phenomenon and the high correlations of higher visual areas (ex. LOC) to this matrix indicates consistency with previously reported findings (Kriegeskorte et al., 2008).

If recognition performance is key to driving correspondence between model and brain representations, then improving model recognition performance should also improve correlations between model layers and visual areas. We found that the model’s correlations increased as a function of its optimization on ImageNet (Figure 4). For each step the model took toward better performance, it also became increasingly similar to neural data. As is known from previous work (Kay et al., 2008; Dumoulin and Wandell, 2008), spatial receptive fields (pooling of inputs) plays a significant role in voxel responses of early vision. We also observe this — Layers 1 and 2 have higher RDM correlations with V1 than Layer 3 even before the model has been highly optimized. However, the pooling structure of our model alone cannot explain these results since as the model becomes optimized, its similarity to V1 and other areas increases, despite the pooling of the model remaining fixed. LOC is not best explained by Layer 3 until the model has been well-optimized–that is, optimization drives Layer 3 above Layers 1 and 2.

We additionally analyzed intermediate convolutional and normalization stages (Figure 6) by computing their object-averaged RDM correlations to each of the visual areas. We observed that the intermediate convolutional and normalization stages roughly fall between the pooling layers in terms of their mapping to each visual area. For practical reasons, Figure 6 presents the unweighted RDM correlations. Empirically, we observed that randomly selecting 1000 features is insufficient to produce stable RDMs from these model stages. Therefore, we present the unweighted RDM correlations using all of the feature dimensions for each layer because computing many more than 1000 feature weightings was infeasible. This change was necessary because the convolutional and normalization stages contain four to nearly ten times more feature dimensions than the pooling layers. Because we did not utilize feature re-weighting, we were able to reliably estimate noise ceilings for these correlations. Determining noise ceiling for correlations where we used feature re-weighting (Figures 2 and 4) was infeasible because it requires estimating the weights on smaller subsets of the data for which we were unable to learn stable weightings.

## Discussion

By analyzing human BOLD responses to hundreds of images, we were able to compare representations of our deep convolutional neural network to those of early, intermediate, and late visual areas simultaneously, thus extending previous work (Yamins et al., 2014; Cadieu et al., 2014) both to humans and to the hierarchy of topographically and functionally localized visual areas (c.f. Khaligh-Razavi and Kriegeskorte (2014); Güçlü and van Gerven (2015)). We found that a deep convolutional neural network optimized for object recognition had representational similarity to human ventral stream visual areas in a hierarchically consistent fashion — early layers best predicted early visual areas and later layers best predicted later areas. The intermediate convolutional and normalization layers residing between the pooling layers exhibited similar, but not as precisely ordered mapping (ex. the second convolution and normalization layers produce very similar RDMs; Figure 6B). The hierarchical correspondence between the network pooling layers and human cortical visual areas increased as the model’s recognition performance was optimized to perform object recognition, suggesting that the functional constraint of object recognition performance was a key component for representations to emerge that resemble ventral visual stream representations. Taken together, our results suggest that biologically plausible computations (convolution, threshold non-linearities, pooling and normalization, (Carandini and Heeger, 2012)) coupled with the top-level constraint of image recognition performance is sufficient to produce hierarchical representations similar to those found in the human visual cortex.

Our analysis of visual representations averaged BOLD responses and model representations to the same object shown from different views, thus stressing object properties common to different viewpoints over ones that are different between viewpoints. Examining responses to individual exemplar images with a single viewpoint (Khaligh-Razavi and Kriegeskorte, 2014) might give insight into the development of tolerant representations, however, our stimulus set did not include enough repeats of the same image to allow for split-half reliability sufficient to analyze without averaging across all views of an object. A potential concern with our object-averaging procedure is that it might artificially favor stronger representational correspondence between the model and more view tolerant cortical areas (Ito et al., 1995; Rust and DiCarlo, 2010; DiCarlo et al., 2012). However, we did not find this to be the case. Instead, correlations were of comparable magnitude across V1 to LOC to the model, what differed was which layer best correlated with each area. We note that correspondence after object-averaging does not necessarily mean that all visual areas or model layers have highly tolerant representations; incidental properties of objects that are still not averaged out across different views might also drive correlations between the model and cortical responses.

The notion that the visual system is hierarchically organized (Felleman and Van Essen, 1991) suggests that intermediate visual areas like V2 and V3 contain intermediate representations, but intermediate in what sense? Our results demonstrate that similar intermediate representations naturally emerge from a deep convolutional network as object recognition performance is optimized, suggesting that the top-level object recognition constraint is sufficient to constrain these intermediate representations. An alternative is that similarity to intermediate areas might only emerge when each model layers are independently optimized for a relevant task (e.g. edge detection for the first model layer, curvature conjunctions for middle layers, object recognition for the higher layers). Nothing in the training of the neural network forced representations to conform to the intermediate representations in visual cortex - the neural network could have learned to generate categorical representations through completely different intermediate mechanisms than those in visual cortex, but our evidence suggests otherwise. While our approach differs from others who have sought to understand what explicit computations might be done in intermediate areas, such as for curvature (Pasupathy and Connor, 2001; Pasupathy and Connor, 2002; Sharpee et al., 2013), angles (Ito and Komatsu, 2004) or for conjunctions of orientations (Anzai et al., 1999; Gallant et al., 1993; Hegde and Van Essen, 2006; Anzai et al., 2007) or other features (Gallant et al., 1996; Freeman et al., 2013), we note that our results do not preclude such an understanding of intermediate visual areas. To what extent the intermediate layers of the model can be characterized as making such explicit computations is a matter of continued investigation which could, in principle, be done by analysis of the receptive field properties of neural network units (Zeiler and Fergus, 2014).

We found that training on object recognition performance was sufficient to drive representational similarity between the model and the human visual system suggesting that model performance and not the specific model architecture was the important factor. Indeed previous analyses (Yamins et al., 2014) showed the strongest correlations of model to monkey physiology data was driven by object recognition performance rather than any specific model parameter. This suggests that the exact formulation of the neurally inspired operations (Carandini et al., 1997; Carandini et al., 2005) like convolution (linear RFs), rectification (spike threshold), normalization and pooling in a layered architecture are less important than the top-level object recognition constraint. Therefore, if other hierarchical models of visual processing similar in architecture to ours, such as HMAX (Riesenhuber and Poggio, 1999), could be trained to have much higher recognition performance, its representations might become more predictive of the hierarchy of the human ventral stream.

While we trained our network solely for object recognition, human visual areas including in the ventral stream likely subserve a multitude of visual functions, and moreover some make connections into dorsal stream areas in parietal cortex thought to subserve other functions such as action planning (Goodale and Milner, 1992). Why then is object recognition performance sufficient to create representations in our model similar to the visual cortex? Object recognition performance may instantiate representations that also support read-outs for other object properties such as position or 3D orientation that might be important for visual functions such as action planning. Alternatively, but not mutually-exclusively, training networks to perform multiple different tasks may better constrain representations to match across multiple visual areas in humans.

There are many facets of visual representation in human visual cortex which are not adequately predicted by the model. For instance, we found that the animate-inanimate distinction was a better predictor of higher visual area responses than our model. However, animacy, like many high-level semantic categories (Kanwisher et al., 1997; Steeves et al., 2006; Epstein and Kanwisher, 1998; Konkle and Oliva, 2012) is not (yet) image-computable and therefore does not represent a model of visual processing. It may be that top-down input representing linguistic, semantic and other cognitive factors or high-level conjunctions and associations between complex stimuli may be required to fully explain high-level representations, particularly for complex representations in the most anterior parts of the ventral stream (Murray et al., 2007). It may also be the case that additional task constraints such as better model training performed on even more realistic object-recognition challenges than ImageNet categorization (Yamins et al., 2014) are needed to improve the model correspondence to human visual areas. Our model also does not yet predict the discrete changes in representation for successive and neighboring areas in human visual cortex for low-level visual features like decrements in image contrast (Gardner et al., 2005) or motion coherence (Costagli et al., 2014). Nor does it predict spatially compact clusters of similar representations such as those found in face patches (Kanwisher et al., 1997; Freiwald and Tsao, 2010). Human object and, in particular, face recognition display particular phenomenology (Tanaka and Farah, 1993; Yin, 1969; De Haan et al., 1987) that may be different from the phenomenology and the types of errors that are made by deep convolutional networks. Thus suggesting that deep convolutional networks using current training regimes are not recapitulating all aspects of human object vision and representation (Szegedy et al., 2013; Nguyen et al., 2014). Nonetheless, our results here suggests that starting with biologically inspired computations and a top-level description of just one important function of the visual system can provide a sufficient starting point for explaining representations in the whole set of hierarchical visual areas we examined in human cortex. Our results thus challenge the idea that each visual area in the hierarchy of visual areas should be understood as having a cicumscribed and easily definable function.

